# Two ways to learn in visuomotor adaptation

**DOI:** 10.1101/2024.11.02.621678

**Authors:** Yifan Zhang, Sana Jayaswal, Nicolas Schweighofer

## Abstract

Previous research has demonstrated significant inter-individual variability in the recruitment of the fast-explicit and slow-implicit processes during motor adaptation. In addition, we previously identified qualitative individual differences in adaptation linked to the formation and updating of new memory processes. Here, we investigated quantitative and qualitative differences in visuomotor adaptation with a design incorporating repeated learning and forgetting blocks, allowing for precise estimation of individual learning and forgetting rates in fast-slow adaptation models. Participants engaged in a two-day online visuomotor adaptation task. They first adapted to a 30-degree perturbation to eight targets in three blocks separated by short blocks of no feedback trials. Approximately 24 hours later, they performed a no-feedback retention block and a relearning block. We clustered the participants into strong and weak learners based on adaptation levels at the end of day one and fitted a fast-slow system to the adaptation data. Strong learners exhibited a strong negative correlation between the estimated slow and fast processes, which predicted 24-hour retention and savings, respectively, supporting the engagement of a fast-slow system. The pronounced individual differences in the recruitment of the two processes were attributed to wide ranges of estimated learning rates. Conversely, weak learners exhibited a positive correlation between the two estimated processes, as well as retention but no savings, supporting the engagement of a single slow system. Finally, both during baseline and adaptation, reaction times were shorter for weak learners. Our findings thus revealed two distinct ways to learn in visuomotor adaptation and highlight the necessity of considering both quantitative and qualitative individual differences in studies of motor learning.

## Introduction

Motor adaptation, a type of motor learning that allows return to baseline performance by reducing the error caused by an external perturbation, is generally thought to depend on two main synergistic systems, e.g., (Huberdeau et al., 2015; Lee & Schweighofer, 2009; Mazzoni & Krakauer, 2006; Smith et al., 2006; Taylor et al., 2014). The first system exhibits fast learning and short-term retention of prior learning, necessitates extended preparation times for expression, and is thought to operate explicitly. Conversely, the second system exhibits slow learning, remains temporally stable during short intervals, can be expressed at short reaction times, and operates implicitly.

Significant advances in understanding these two systems have been possible via a combined experimental and modeling approach. The dual-rate model (Smith et al., 2006) for a single task, and its extension to multiple tasks (Lee & Schweighofer, 2009), accounts for several behavioral phenomena, such as anterograde interference, spontaneous recovery, rapid unlearning, and some form of savings. In these models, the fast and slow processes compete for the same error, yielding the prediction that the fast-learning-fast-decaying process decays later in learning when the remaining error becomes small. At the population level, the slow process has been shown to predict long-term retention (Joiner & Smith, 2008). A recent study (Hadjiosif et al., 2023) further delineated the different contributions of a temporally volatile process and a persistent learning process in predicting savings and long-term retention, respectively, with the processes tentatively mapping onto a fast and a slow process.

It has long been recognized that motor learning exhibits considerable individual variability, e.g. (Ackerman, 1988; Keele & Hawkins, 1982), recent work has begun to study individual variation in motor adaptation, e.g., (Della-Maggiore et al., 2009; Miyamoto et al., 2020; Stark-Inbar et al., 2017; Trewartha et al., 2014). For instance, (Miyamoto et al., 2020) showed that learners varied widely in how the implicit and explicit processes sum to the total adaptation level, with some learners displaying high implicit but low explicit learning and vice versa. Recruitment of the explicit process, which has been linked to spatial working memory (Anguera et al., 2010; Vandevoorde & Orban de Xivry, 2020), involves increased reaction times (Haith et al., 2015; McDougle & Taylor, 2019). Accordingly, learners who show faster learning and smaller aftereffects associated with the explicit process exhibit longer reaction times (Fernandez-Ruiz et al., 2011).

Whereas these previous studies proposed quantitative differences in the recruitment of fast and slow processes in adaptation, we (Oh & Schweighofer, 2019) recently exposed qualitative differences in adaptation to a 20-degree visuomotor rotation, with two classes of learners: those who only updated a single “baseline” or (“body” (Berniker & Kording, 2011)) process showed gradual de-adaptation and little or no savings, and those who additionally created a new memory showed savings and a quick return to baseline in the absence of perturbation.

Here, we, therefore, hypothesized that learners in a motor adaptation experiment with a 30- degree rotation would belong to one of two qualitatively different groups: strong learners, who engage the competing fast and slow processes, and weak learners, who only engage the slow process. The strong learners are expected to show large variability in recruitment of the competing fast and slow processes. In addition, we also posit that in strong learners, the slow process would predict 24-hour retention and the fast process would predict savings, as recently suggested by (Hadjiosif et al., 2023). In weak learners, the process would be slow and only predict retention. Finally, we test whether weak learners exhibit shorter reaction times than strong learners.

## Methods

### Participants

We recruited 44 participants (21 Females, 23 Males) for a 2-day experiment via the *Prolific.com* online recruitment platform designed for online scientific research studies (Palan & Schitter, 2018). Participants were residents of the United States, and reported to be fluent in English, 19 to 40 years old (29.1 ± 4.1; all results as mean ± standard deviation unless noted) and right-handed. Participants received monetary compensation of $10 per hour and a $5 completion bonus if they completed the second session in a 24 plus or minus 5 hours window. Informed consent was obtained before the study began, and the Institutional Review Board of the University of Southern California approved all procedures.

### Experiment Setup

The online experiment was modified from (Tsay et al., 2021). The task required participants to shoot at targets positioned at eight pseudo-random locations on a circle. On perturbation trials, a counterclockwise visuomotor rotation of 30 degrees was applied to the cursor (**Figure** 1 A). At the start of each trial, participants positioned their cursor on a centered fixed point corresponding to the home position. A target appeared 500 ms later at one of the 8 locations in pseudo-random order. Participants were instructed to aim directly at the target by moving their index fingers on a laptop trackpad (**Figure** 1 B). If the duration of the cursor movement from the home position to the target circle exceeded 1 second, a “too slow” signal was displayed. In data analysis, however, all trials are considered valid regardless of movement time.

**Figure 1.**
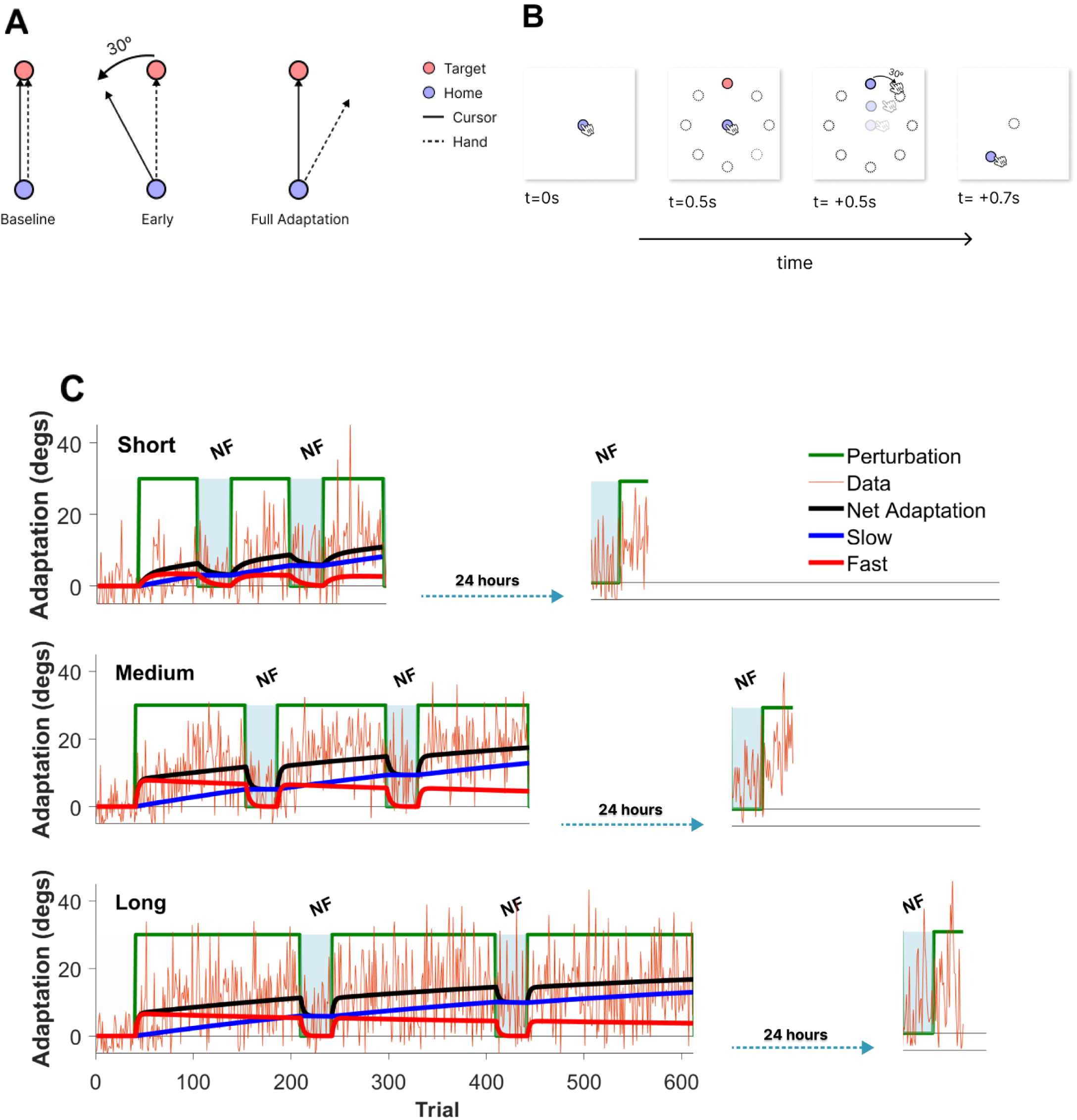
The visuomotor adaptation experiment. **A.** Illustration of the adaptation task, in which a 30-degree counterclockwise rotation was applied to the cursor as participants reached a target. To counter this perturbation, subjects needed to learn to shoot at 30 degrees to the right of the target by moving their index finger on a laptop trackpad. **B.** Timeline of each trial during the adaptation phase. First, a fixation point (solid blue) representing the home position appeared in the middle of the screen. Participants moved the cursor to the fixation point to start the trial. After a 0.5s delay, one of 8 targets (separated by 45 degrees, dotted circles) appeared pseudo-randomly on the target circle. During adaptation trials, the cursor trajectory, rotated 30 degrees relative to the finger position, was shown in real-time (solid circle), and the endpoint position feedback was shown for 0.2s before starting the next trial. “+ … s” indicates the minimum time needed to complete each step of a trial. During no-feedback trials, neither the trajectory nor the endpoint position was shown. **C.** Experiment schedules and examples of the fast/slow model fit for three subjects, one for each dose level. Day 1 sessions were separated from day 2 sessions by approximately 24 hours. The green lines indicate the rotation in degrees. “NF” indicates the no- feedback trial blocks. On day 2, the first no-feedback block of 32 trials is the retention block, and the second perturbation block of 32 trials is the relearning block. Note the large between-subject variability in adaptation and the diversity in the dynamics of the fast and slow processes in these three examples.

The experiment involved two sessions, with the second occurring within 24 ± 5 hours after the first session. On day 1, following instructions, which contained a short video, and a questionnaire about demographic information, participants performed a familiarization block of 16 trials, followed by a baseline block of 40 trials. Participants were randomly assigned to one of three training schedules based on a small, medium, or large dose of perturbation trials. Each group performed three training blocks, with each block separated by a 32-trial no-feedback block (**Figure** 1 C). The small dose group (n=16) completed 56 trials per block, the medium dose group (n=15) 112 trials, and the large dose group (n=13) 168 trials.

During feedback trials, both the cursor’s online position was displayed in real-time. The final position was then shown for 0.2s where the cursor intersected the target circle. During no-feedback trials, participants were only shown the targets and were instructed to aim directly at the targets. The second session on day 2 (**Figure** 1 C, rightmost), which was identical for all participants, started with a no-feedback block of 32 trials to assess 24-hour retention, then a relearning block of 32 trials to assess 24-hour savings.

### Data Analysis

Data were downloaded from Firebase as JavaScript Objective Notation files (JSON) and converted into csv files. At each trial, the adaptation level was computed as the angle between the position where the cursor crossed the target circle minus the target angle. Trials with an angle greater than 60 degrees or less than -30 degrees were classified as outliers. We replaced outliers with the angle means of their 8 non-outlier neighbors. The total percentage of outliers was 1.6% for the overall data and 3.2% or less for any individual data.

In the session of day 2, we measured the retention level by computing the mean of the adaptation level of the no feedback 32-trial retention block. To measure savings, we removed any bias by subtracting the retention level from the mean adaptation in the 32 relearning trials. We then computed savings as the difference between this baseline-corrected relearning measure and the mean initial adaptation in the first 32 trials of the first learning block on day 1.

Reaction times (RTs) were extracted from the raw data and computed as the time between the presentation of the target and the cursor crossing the 1 cm circle from the start position, as in (Tsay et al., 2021). We computed the median RT both in baseline trials and in training trials when the perturbation was 30 degrees.

### Model-based analyses

#### Fast/Slow Adaptation Model

We fitted a noiseless parallel two-state dual rates model (Smith et al., 2006) to the day 1 adaptation data of each subject. This model captures both a fast *xf* state and a slow *xs* state, each corresponding to a motor memory that sums up to give the estimated adaptation level (in degrees). The error between the perturbation amplitude and the model output updates both states. For each trial *t* and subject *i*, the model is thus described by the state update equations for the fast and slow processes, the error equation, and the output equation:

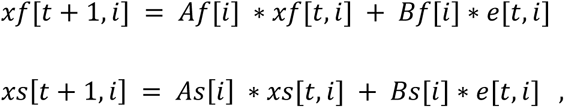

where the error *e*[*t*, *i*] is given by:

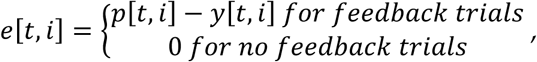

and the output is given by the sum of the fast and slow processes:

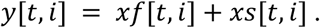

The learning rates are constrained as *Bf* > *Bs* > 0 and the forgetting rates as *As* > *As* > 0 for all s, to ensure faster learning and forgetting of the fast process. Note that during no feedback trials, both states decay exponentially. In additional analyses, we fitted the data to a single-process model, similar to the dual-process model above but with a single state.

#### Fitting the models to the reaching angle data

Processed data were analyzed using MATLAB (The MathWorks). We used the *fmincon* global optimization function with the interior point method to minimize the mean squared error between the model-predicted angle and actual angles. Two states and one state models were individually fitted to each subject’s day 1 data. The learning rates were constrained between 0 and 0.5 to prevent excessive oscillations. To enhance the robustness of the fitting process and to avoid values close to 1, we re-parameterized the forgetting rates into their time-constant equivalents, mapping the range [0, 1] to positive real numbers- see (Kim et al., 2015), and used the time constants as free parameters. For instance, for the two-state model,

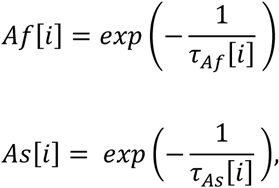

where *τ_As_*> *τ_Af_*> 0 are the time constants for the slow and fast processes. We used a loss function that equally weighted feedback and no-feedback trials to adequately fit the decay during no-feedback trials and ensure similar fit results across doses. Furthermore, we initialized the model with 500 distinct initial parameter conditions to avoid local minima. We analyzed the estimates of slow, fast, and total adaptation at the end of training by extracting these values on the last trial of day 1.

#### Parameter recovery simulations

We designed our experiment to improve the estimation of the fast and slow process at the individual level. Most previous adaptation studies included a single learning block, a decay (or washout) block, and a relearning block, e.g., (Joiner & Smith, 2008; Smith et al., 2006; Taylor et al., 2014). Because parameter estimation of the fast-slow model is relatively poor in such single-block designs, previous studies fitted the model to average data, improving fit because of reduced trial-by-trial variability. Here, to improve the fit of the models to individual data, we designed our experiment with three learning blocks, in which the fast and slow processes are updated via errors, and two short retention blocks, in which the fast process quickly decreases. To show the advantage of this multi-block design in parameter estimation, we performed parameter recovery in simulations and compared the recovered parameters in our design and a design with a single learning block. We simulated patients using one- and two-process models with realistic trial-by-trial variability. True parameters (n = 44) were sampled from estimated model fit parameters to actual data with replacement. We added zero-mean Gaussian noise to the model output first using a standard deviation of 4.4 degrees, which corresponds to the average standard deviation in baseline trials in the experiment, and then using a standard deviation of 6.5 degrees, which is ∼50% above the baseline average. The schedule for each simulated subject in the three-block experiment was randomly chosen among the long, medium, and short schedules with the same proportion as in the experiment. The schedules for the simulated single-block experiment contained the same number of learning and decay trials as in the three-block experiment but concatenated in one adaptation block followed by one forgetting block. We then fitted the simulated data with a corresponding one or two-process model. The recovery error for each parameter was computed as the mean relative error ((recovered-true)/true) in percentages.

#### Predicting the adaptation from baseline reaction times

Because we observed lower baseline reaction times in weak learners, we tested how well median baseline reaction time could predict subjects as strong or weak learners. For this, we used a weighted logistic regression with inverse frequency weighting. Weighted logistic regression (Lee & Liu, 2003) is an extension of logistic regression that accounts for class imbalance by assigning different weights to classes based on their prevalence (13 weak learners and 31 strong learners in our case). For each subject i, the weighted logistic regression model predicts the probability of being a strong learner as:

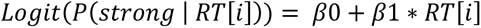

Where *Logit* is the log odds and *β*0 and *β*1 are the intercept and slope estimated during model fitting. To validate the robustness of the model, we employed a standard bootstrapping procedure, which involves repeatedly resampling the data with replacement and refitting the model to each resampled dataset with 1000 bootstrapped samples.

## Results

### Cohort-level analysis and modeling

We first analyzed data from all subjects by assuming that all learners develop explicit and implicit processes, as modeled by fast and slow states. The model fit was good overall, with the RMSE between the model and data of 7.7 + 1.5 degrees, with a 95% CI range of 4.3 – 10.6 (compared to the baseline mean standard deviation of 4.4 degrees, with a 95% CI range of 1.9 - 9.2). Parameters show a wide range of dispersion in estimated parameters, indicating high between-subject variability (*Af*: 0.68 ± 0.2 (mean ± SD); *As*: 0.99 ± 0.11; *Bf*: 0.21 ± 0.20; *Bs*: 0.01 ± 0.01).

Thanks to our design with three learning blocks, parameter recovery was good overall, as shown by the small recovery error between the true (known) and estimated parameters (see Supplementary Figure S1). The recovery performance with three learning blocks largely surpassed that with a single learning block, especially for the time constant of the slow process (see Supplementary Figure S2 for details).

#### Slow and Fast Processes predict retention and savings, respectively

We then tested whether the fast and slow processes at the end of day 1 predicted the 24-hour savings and retention, respectively. This hypothesis is based on three previous studies. A first study showed that when a fast/slow process was fitted to average data in multiple dose groups, the slow process predicted 24-hour retention (Joiner & Smith, 2008). Next, we previously showed that the slow implicit process accounted for retention at the individual level in a 10-minute retention test (Lee et al., 2018). Finally, a recent study showed that a component of motor memory that is temporally persistent after 60 seconds contributes to long-term retention, whereas a temporally volatile component that has decayed by 60 seconds contributes to savings (Hadjiosif et al., 2023).

As hypothesized, the slow process at the end of day 1 training positively correlated with retention (R = 0.57, p = 0.00005, **Figure** 2 A), and the fast process positively correlated with savings (R = 0.49, p = 0.0007, **Figure** 2 D). Conversely, the fast process showed a non-significant correlation with retention (R = -0.21, p = 0.18, **Figure** 2 C), and the slow process showed a non-significant correlation with savings (R = -0.27, p = 0.08, **Figure** 2 B). (Note that the moderate negative correlation between slow process and savings and fast process with retention are expected because the slow and fast processes are negatively correlated - see below and **Figure** 3 A).

**Figure 2.**
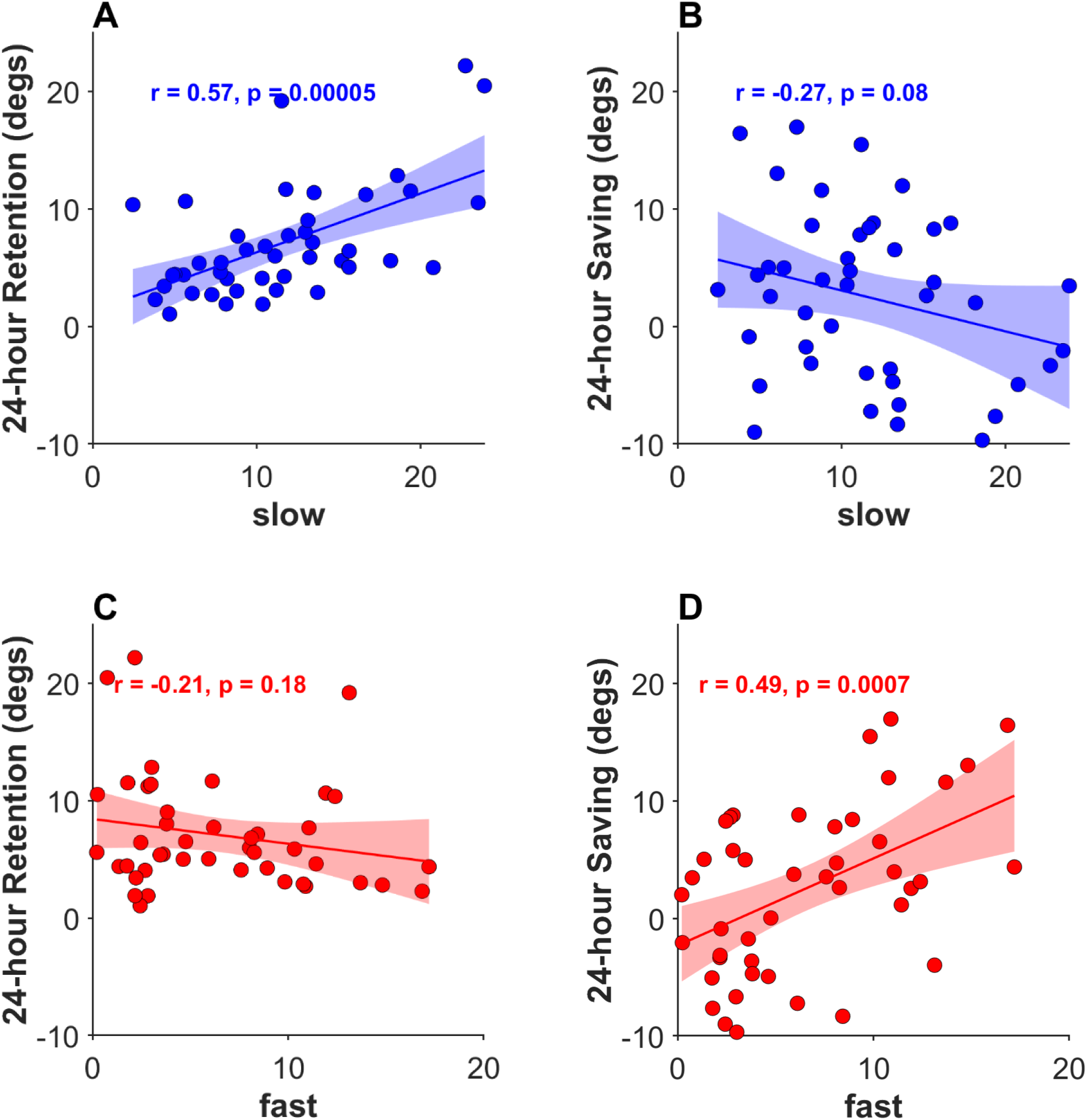
Predicting 24-hour retention and savings from the fast and slow model at the end of the training session on day 1. **A.** Strong positive correlation between the slow process and retention. **B.** Non-significant negative correlation between the slow process and saving. **C**. Non-significant correlation between the fast process and retention. **D**. Strong positive correlation between the fast process and saving. The shaded areas show the 95% confidence interval of the population mean. Each dot represents a subject.

**Figure 3.**
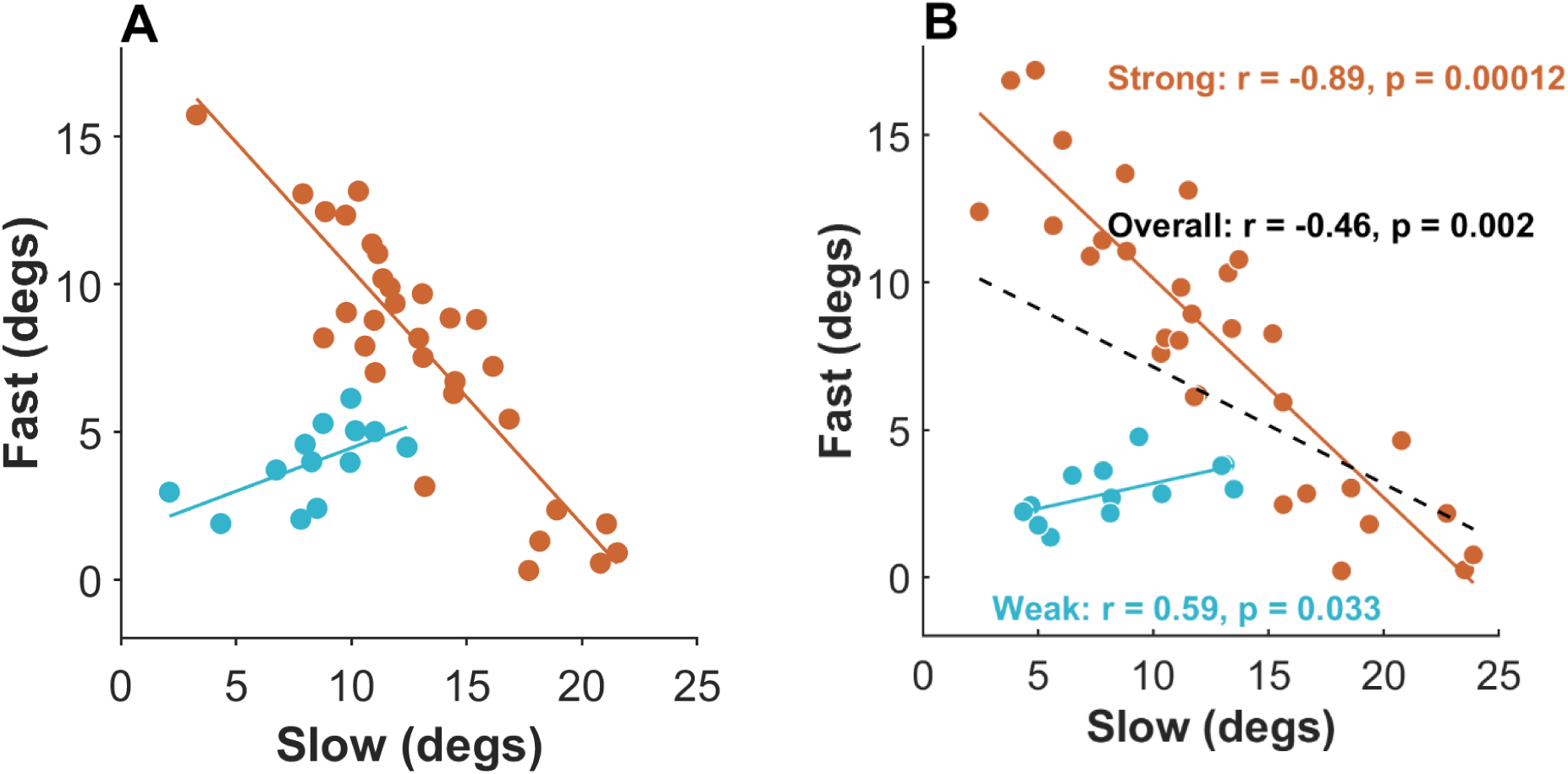
Relationships between fast and slow processes at the end of day 1 training for both strong and weak learners in simulations and the experiment. **A**. Fit of two state models to simulated data generated by two state models with noise added to the model’s output for 44 simulated subjects. The red group (31 subjects) represents fast and slow processes from a 2-state model at the end of day 1, where the forgetting rates (*As* and *Af*) are fixed, and the learning rates (*Bs* and *Bf*) are sampled from normal distributions. The cyan group (13 subjects) illustrates two processes estimated from simulations of a single-state model. For this group, we first simulated 44 subjects using single-state model, where the forgetting rate (*A*) is fixed, and the learning rate (*B*) is sampled from a normal distribution. Then, to match the characteristics of actual weak learners, the 24 simulated subjects with one process near zero (defined by 0.3 or lower for the recovered *Af* parameter) were excluded, and the additional 13 subjects were randomly sampled from the remaining 20 subjects. **B.** Fit of two state models to actual data. Correlation scatter plots and regression lines between fast and slow processes at the end of training in day 1 estimated from data for each subject. Colors show group identity based on k-mean clustering. Red: strong learners; cyan: weak learners. Black line: regression line for all learners.

#### Individual variability washes out the effect of dose

Whereas we designed our experiments along that of (Joiner & Smith, 2008) with three doses of training with a predicted effect of dose on retention, the large inter-individual variability in adaptation in our data washed out any effect of dose. One-way ANOVAs across all doses yielded non-significant results for total adaptation at the end of training (average of adaptation over last 16 trials, p = 0.54), retention (p = 0.13), and savings (p = 0.75). Additionally, linear models (*Retention ∼ Slow *Dose*, and *Savings ∼ Fast *Dose*) with dose level as a factor and with the short dose as the reference showed a non-significant effect of dose on the slow process with retention (p = 0.32 for the medium dose and p = 0.56 for the long dose compared to the short dose) and the fast process with savings (p = 0.29 for the medium dose and p = 0.74 for the long dose compared to the short dose). Thus, despite large differences in the number of adaptation trials between the three doses (56*3=168, 112*3=336, 168*3=504 trials), the large inter-individual variability washed out the effect of doses. As we will see in the following, the variability is due to both qualitative and quantitative differences between subjects.

#### Analysis of Individual Differences in Learning, Retention, and Savings

Given the large variability in performance at the end of day 1, and our previous study suggesting two qualitatively distinct groups of learners (Oh & Schweighofer, 2019), we used k-means clustering with k=2 to classify learners across all doses into two learnability levels based on the mean of the last 8 trials of day 1. This clustering resulted in 31 strong learners and 13 weak learners, with a cut-off of 15.0 degrees (Supplementary Figure S3).

Strong learners showed significant retention and savings (one-sample t-test, p = 0.0001, 7.8 ± 5.2 degrees for retention; one-sample t-test, p = 0.0056, 3.9 ± 7.3 degrees for saving). In contrast, weak learners exhibited significant retention (one-sample t-test, p = 0.0001, 5.3 ± 3.0), but no savings (one-sample t-test, p = 0.59, 0.8 ± 5.4). Accordingly, there was a difference in savings between strong and weak learners (2-sample t-test, p = 0.041), but no difference in retention (2-sample t-test, p = 0.096).

#### Compensation between fast and slow processes in strong but not in weak learners

The above findings for strong and weak learners suggest that 1) strong learners rely on two processes that predict retention and savings, and 2) weak learners rely on a single memory process that only predicts retention. We now provide additional evidence for these insights in four complementary model-based analyses.

First, a BIC analysis of two- and one-state models to the data supports the recruitment of two processes in the strong-learner group and a single process in the weak-learner group (Supplementary Figure S4 A, B). The bootstrapped BIC distributions showed that, for the strong learner group, the BIC confidence intervals are much lower for the two-state model than the one-state model. In contrast, for the weak learner group, the BIC confidence intervals overlap for the two models, supporting the simplest, one-state model.

Second, the two groups showed qualitative differences when we plotted the slow state as a function of the fast process estimated at the end of day 1. Because the fast and slow components of the model sum up to the total adaptation, and assuming approximately equal adaptation *y*[*t_end_*] at the end of adaptation *t*_end_, the fast and slow states are negatively correlated, with a slope of -1 (because *x*[*t*_end_, *i*] = − *x*[*t*_end_, *i*] + *y*[*t*_end_]; see model equation above, and simulations in Figure 3A). Correspondingly, the correlation between slow and fast processes at the end of day 1 for all 44 subjects in the experiment was negative (r = -0.46, p = 0.002) (Figure 3 B black).

However, the correlation patterns differ when separating the subjects according to learnability. For strong learners (Figure 3 B red), the slow and fast processes showed a strong negative correlation (r= -0.89, p = 0.00012), and wide dispersion along this line (standard deviation: 4.8 degrees), due to largely variable learning rates (*Bf_strong_*: 0.3 ± 0.21, *Bs_strong_*: 0.01 ± 0.01). Such dispersion along the negative slope can be seen in simulations of two-process models in which the learning rates were independently sampled from normal distributions capped between 0 and 1 (*Bf* = 0.3 ± 0.21, mean *Bs* = 0.01 ± 0.01, with fixed retention rates *Af* = 0.75, *As* = 0.95; **Figure** 3 A red).

In contrast, weak learners (**Figure** 3 B cyan) showed a positive correlation between the two processes (r = 0.59, p = 0.033). A linear model (*Slow ∼ Fast * Learning Level)* further confirmed the significant effect of learning level in modulating the slopes for the two groups (p = 0.0098). The positive correlation for weak learners is due to the estimation of two slow processes: the estimated learning rates (*Bf*: 0.043 ± 0.07; *Bs:* 0.03 ± 0.06) are highly positively correlated (r = 0.78). To verify that this result is compatible with a single underlying process generating the data, we simulated multiple subjects with a single-state model (with *A*: 0.95 ± 0.02; *B*: 0.04 ± 0.2; noise std = 4.4), and then fitted the same 2-states model as above to these simulated data and estimated slow and fast process at the end of training. The resulting regression line has a positive slope, qualitatively matching weak learners in data (**Figure** 3 A cyan; additional simulation details in legend).

Third, we predicted that the savings and retention in weak learner groups are consistent with the single process. Indeed, the estimated single process at the end of day 1 (**Figure** 4 A) correlated with retention (p = 0.001, R = 0.80) but not with savings (**Figure** 4 B, p = 0.422, R = -0.25). In addition, the single time constant of weak learners and the time constant of the slow process of strong learners showed no differences (t-test h = 0, p = 0.461; 95% CI for *As* = [0.94,0.99] in strong learners; 95% CI for *A* = [0.88,0.99] in weak learners).

**Figure 4.**
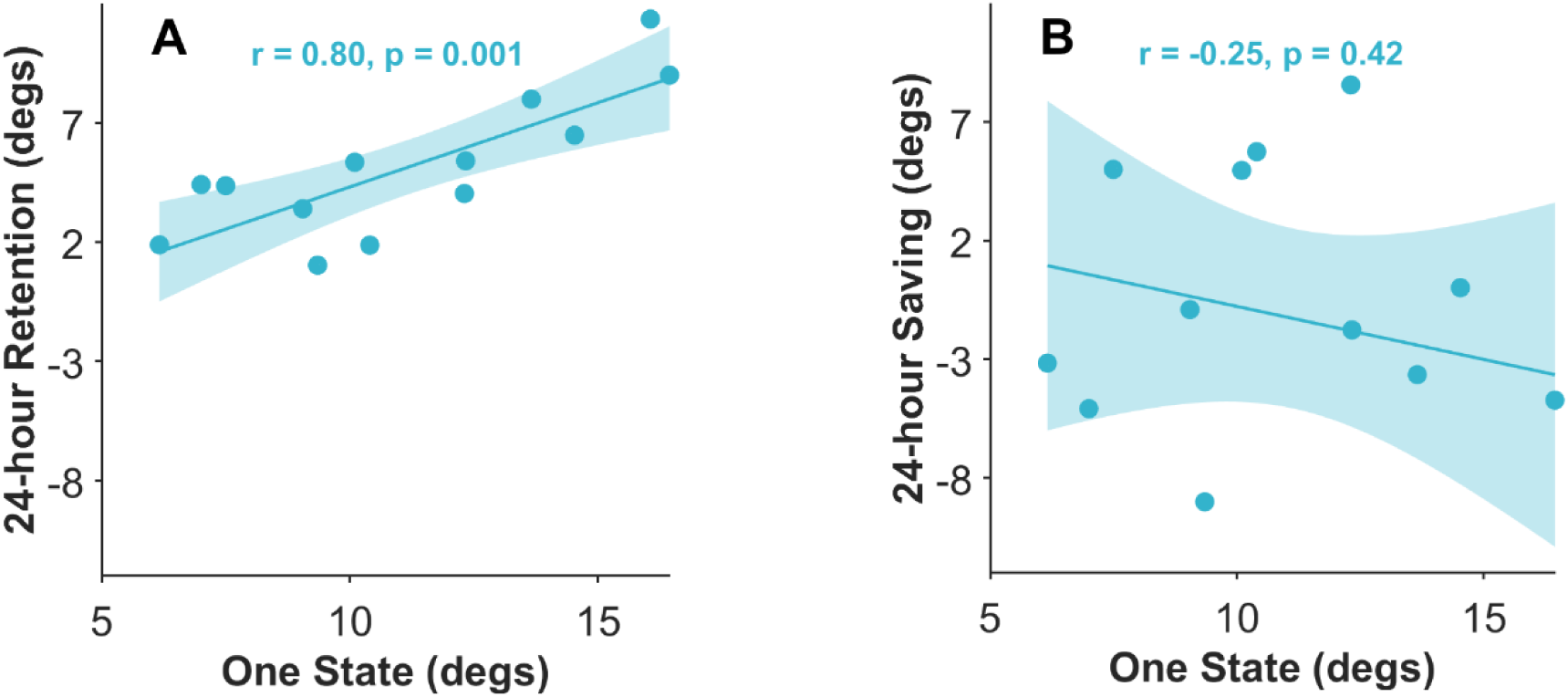
**Predicting 24-hour retention and savings from one state model at the end of day 1** in weak learner’s group. One-state model estimate of adaptation at the end of day 1 vs retention (A) and savings (B). Positive correlation between estimated adaptation at the end of day 1 and retention (R = 0.80, p = 0.001) but not with saving (R = -0.25., p = 0.42)

Finally, we tested for possible differences in RTs between the two groups. Previous studies have shown that longer reaction times are associated with more cognitive/slow processes (Fernandez-Ruiz et al., 2011; Haith et al., 2015; Taylor et al., 2014). We, therefore, predicted shorter reaction times for weak learners during learning. Indeed, during learning trials (**Figure** 5 B), strong learners also exhibited a larger RT (median 442 ms; IQR = [426, 495]) than weak learners (402 ms; IQR = [356, 448]; p = 0.027; Mann–Whitney test). However, surprisingly, during baseline trials (**Figure** 5 A), strong learners also exhibited a larger RT (median RT =443 ms, and IQR = [405, 501]) than weak learners (RT = 384 ms, and IQR = [364, 406]; p = 0.034, Mann–Whitney U test). A linear regression model showed that RT at baseline predicted the level of learning for all subjects (average of adaptation in the 8 last trials of day 1) (0.023, *p* = 0.039). We then evaluated the ability of baseline reaction time (RT) to predict strong and weak learner groups using a weighted logistic regression model. Overall accuracy was 61.4% (compared to 50% accuracy of a baseline classifier), with a 95% bootstrapped CI ranging from 49% to 84%.

**Figure 5.**
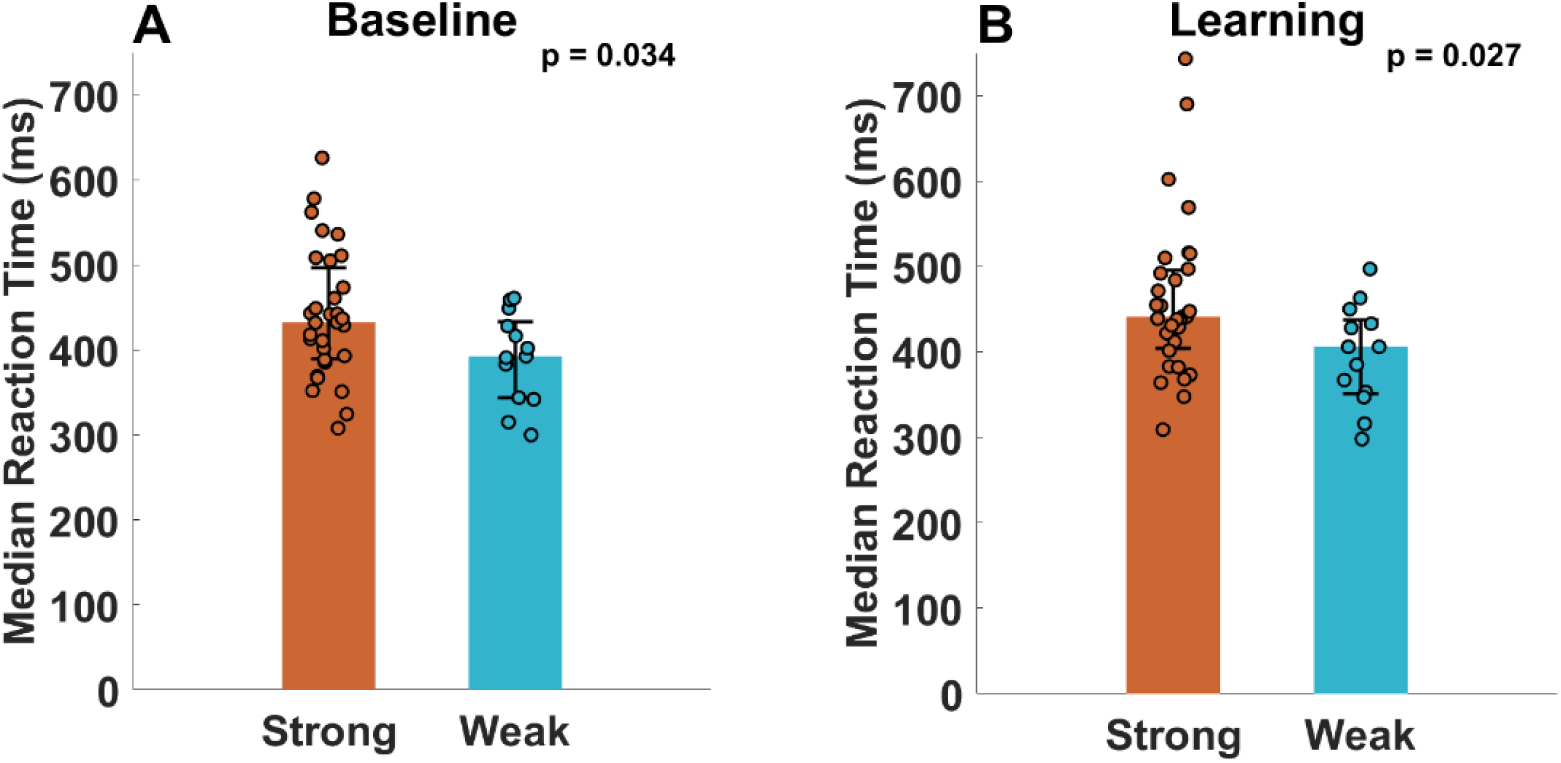
Reaction times are larger in strong learners than weak learners both during baseline (A) and learning (B) trials. Dots: median for each subject. Bar height: the median RT. Error bar: IQR.

## Discussion

Whereas fast and slow models of adaptation are typically fit to data averaged across subjects in single block designs, here we designed an adaptation experiment with three learning two forgetting blocks to estimate the model’s parameters in individual learners. A parameter recovery analysis confirmed that this three-learning block design led to more accurate parameter estimation than a single-learning block design. We then showed that, despite large differences in the number of adaptation trials in the three dose groups, the inter-individual variability overshadowed the expected dose-dependent effects on adaptation, retention, and savings.

The individual variability was both qualitative and quantitative. A clustering analysis identified two qualitatively distinct groups of learners: strong learners, who exhibited robust retention and savings, and weak learners, who demonstrated retention but no savings. Additional model-based analysis showed that strong learners rely on two competitive fast and slow processes, while weak learners rely on a single, slow process. The strong learners showed a large between-subject variability in their recruitment of the fast and slow processes, as shown by a large dispersion along the negative correlation slopes that emerge from the fast and slow models. A recent study found a similarly large between-subject dispersion of implicit and explicit processes along this negative correlation slope (Miyamoto et al., 2020). Furthermore, strong learners exhibited longer reaction times than weak learners during both baseline and learning trials, which further supports that strong learners rely more on cognitive/explicit processes. Finally, larger reaction times during baseline in strong learners suggest a predisposition to engage the cognitive/explicit processes even before adaptation was introduced.

In recent work (Oh & Schweighofer, 2019), we suggested that interindividual differences in the rate of de-adaption and re-adaption to a visuomotor rotation depended on the continuous update of an existing baseline model (resulting in slower de-adaptation and re-adaptation) or on the ability to create and update new internal models (i.e., motor memories) specific to the perturbations, and then easily switch between these models (resulting in fast de-adaptation and re-adaptation). These interindividual differences were controlled by the relative precision of the different memories, which yielded individual differences in model selection and learning rates by modulating “responsibility signals”. This initial model was extended into a new contextual inference model (Heald et al., 2021) that enables detailed, trial-by-trial quantification of how multiple memories are generated, weighted, and updated in response to varying errors and contexts. The link between the contextual learning paradigm and fast-slow models of adaptation warrants further research.

In strong learners, we observed strong relationships between the slow process and retention on one hand and the fast process and savings on the other hand. Thus, our results extend the results of (Joiner & Smith, 2008) to individual retention and savings in a visuomotor adaptation experiment. Our results also suggest that the temporally persistent and volatile memory processes described in (Hadjiosif et al., 2023) are captured by the fast and slow processes, respectively. Note that the reproduction of these prior results with our model-based analysis further validated, at the population level, the goodness of fit of our model to the adaptation data of day 1.

Whereas we modeled only one fast and one slow process, behavioral and imaging experiments have uncovered a third ultra-slow process (Forano & Franklin, 2020; Heuer & Hegele, 2015; Kim et al., 2015; Kording et al., 2007). Because the estimated time constant of the slow process in our study is at most 1000 trials (see Supplementary Figure S1), roughly corresponding to 1 hour, the slow process will have decayed by about 99% in 5 hours. The correlation between the slow process and 24-hour retention suggests that the ultra-slow process is updated based on the slow process, as previously proposed (Criscimagna-Hemminger & Shadmehr, 2008), predicting a large variability in the ultra-slow process related to the variability in the slow process. Additional research with designs that uncover such ultra-slow processes (for instance, via dual-adaptation (Forano & Franklin, 2020)) of multi-day adaptation is needed to test this hypothesis.

The main limitation of our study is the use of an online experiment in a non-controlled environment. In particular, we do not know whether the smaller RTs in weak learners are due to a trait or to possible dual-tasking during the experiment, which is known to reduce adaptation (Taylor & Thoroughman, 2008). In addition, given that online experiment data tends to exhibit higher noise levels than offline data (Tsay et al., 2021), removing online feedback could enhance data quality in online experiments and further improve the model fit (Kasuga et al., 2015).

Despite these limitations, we have successfully reproduced, and extended with a fast-slow model-based analysis, previous results demonstrating i) qualitative differences in adaptation based on whether one or two memory processes are recruited (Oh & Schweighofer, 2019), ii) strong quantitative differences in competitive recruitment of the implicit/fast and explicit slow processes (Miyamoto et al., 2020), at least when these two processes are recruited in strong learners, iii) a double dissociation between slow and fast memories (Hadjiosif et al., 2023), which we extended to a multiple target paradigm, but here again exists only for strong learners, iv) longer reaction times during adaptation associated with involvement of the cognitive/slow processes (Fernandez-Ruiz et al., 2011; Haith et al., 2015; Taylor et al., 2014) in strong learners. In addition, we have shown a novel predictive effect of baseline reaction times, which can predict, to some extent, if a learner will become a strong or weak learner. Thus, our findings underscore the importance of considering both quantitative and qualitative individual differences in motor adaptation. Looking forward, investigating between-subject variability in motor adaptation and long-term retention, ideally in more realistic and relevant 3D tasks (Cesanek et al., 2024; Ferrea et al., 2022), could help design individualized interventions in neurorehabilitation following cortical damage.

## Author contributions

Author contributions: Y.Z. and N.S. designed the research; S.J. performed the experiments; Y.Z. analyzed data; Y.Z. and N.S. interpreted results of experiments; Y.Z. prepared the figures; Y.Z. and N.S. drafted the manuscript; Y.Z. and N.S. edited and revised the manuscript. All authors approved the final version of the manuscript.

