## Supplementary Material for "Two ways to learn in visuomotor adaptation"

### Supplementary Materials

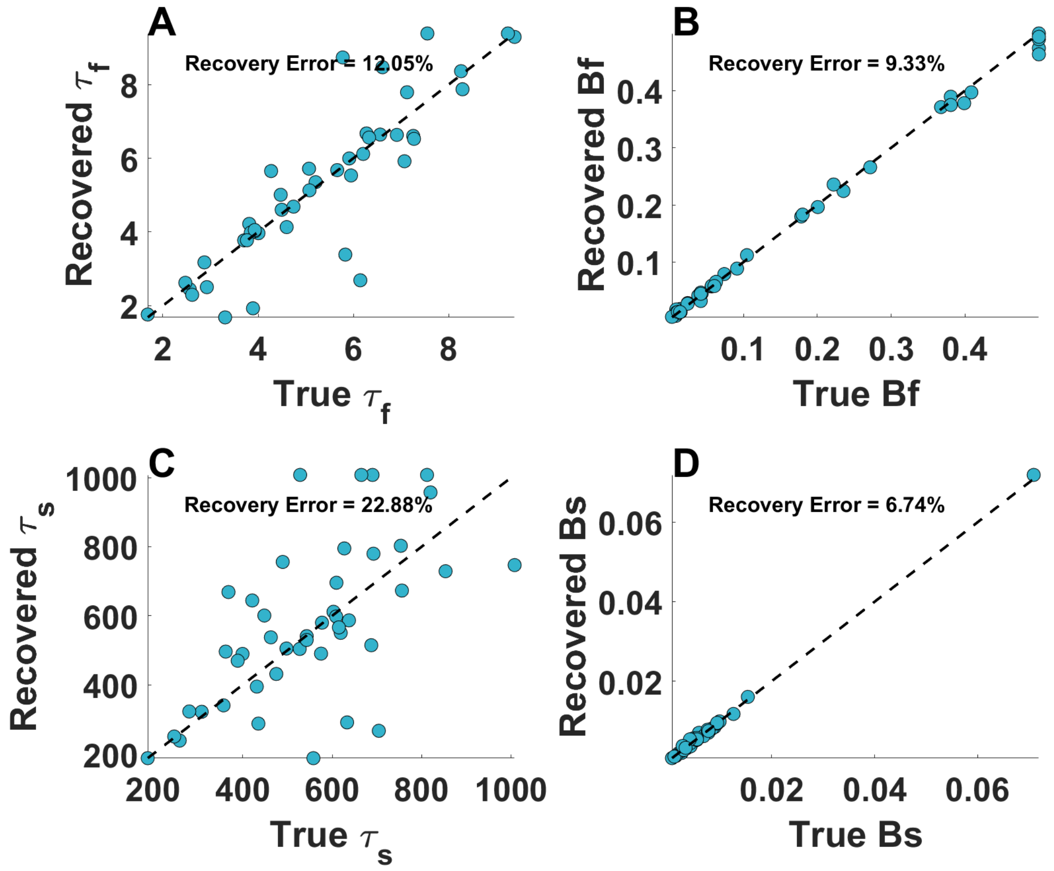


**Figure S1 Parameter recovery error for the two-state model in our adaptation paradigm with three adaptation blocks.** A, B, C, D: the recovery of parameters *Af*, *Bf*, *As*, and *Bs*. Perfectly recovered parameters would fall on the diagonal lines (dashed).

A. Recovery error in one-block and three-block designs

One-state model, same noise level (4.5 degs)

| | *T* | *B* |
|---|---|---|
| 1 block | 38.6% | 15.4% |
| 3 blocks | 15.7% | 9.6% |

B. Two-states model with different noise levels:

| | $\tau s$ | *Bs* | $\tau f$ | *Bf* |
|---|---|---|---|---|
| 1 block<br>Noise: 4.5 degs | 62.2% | 18.1% | 24.9% | 13.3% |
| 3 blocks<br>Noise: 4.5 degs | 22.9% | 6.7% | 12.0% | 9.3% |
| 1 block<br>Noise: 6.4 degs | 70.4% | 36.5% | 45.9% | 32.7% |
| 3 blocks<br>Noise: 6.4 degs | 38.1% | 12.1% | 28.5% | 14.7% |

**Figure S2**, **Comparison of parameter recovery error in simulated experiments with one learning block or three learning blocks with the same number of trials.** A) Comparison of recovery for a one-state model, and B) Comparison of recovery for a two-state model and two levels of noise. Errors were computed as the percentage deviation from true parameters.

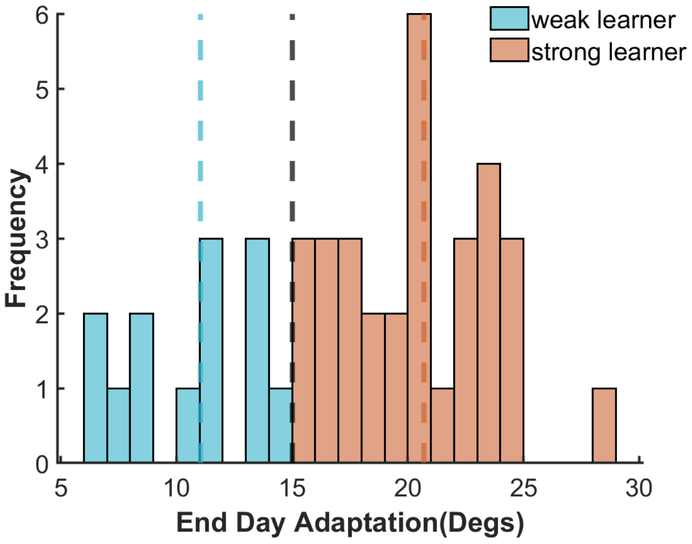


**Figure S3**, **Clustering into weak and strong learners.** K-means clustering (k =2) results for strong (n =31, median = 11.0) and weak (n =13, median = 20.1) learners based on the model-free mean of adaptation in the last 16 trials at the end of day 1. Colored dashed lines mark the median of each population and the black dashed line marks the between-group cut-off (15 degrees).

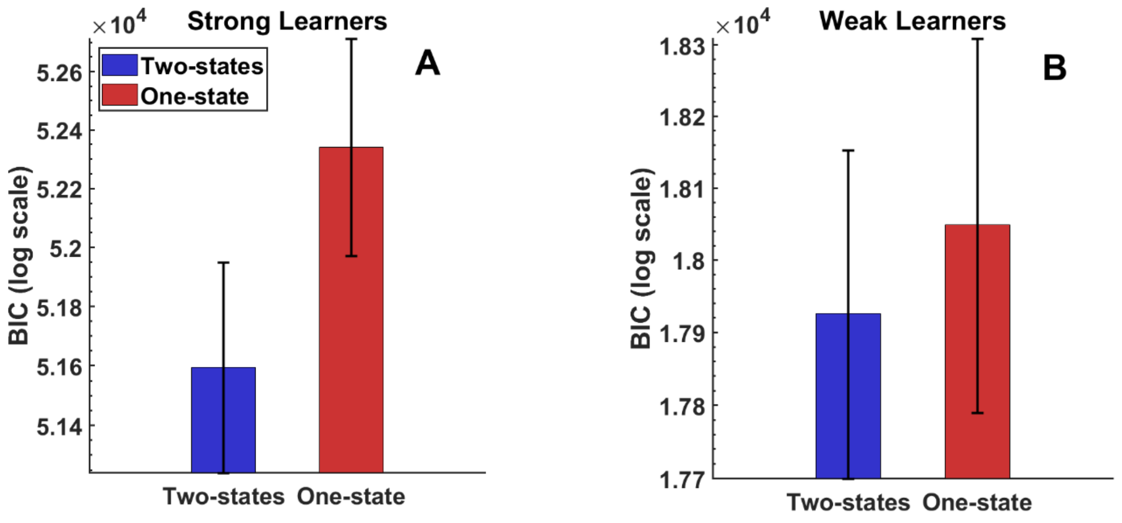


**Figure S4**, **A, B**: **Bootstrapped BIC distributions for strong and weak learners using one- and two-state models**. We fitted the one-state and two-state models to strong and weak learners' groups and compared the 1000 sample bootstrapped BIC distributions between 2 models in each of the strong and weak learner groups. The mean difference was larger for the strong learner (two-state BIC mean = 5.13e+04 ± 212.10 CI [5.13e+04, 5.20e+04]; single-state BIC mean = 5.25e+04 ±189.72 CI [5.20e+04, 5.27e+04 [) than for the weak learner (two-state BIC mean = 1.79e+04 ±107.41 CI [1.77e+04, 1.82e+04]; single-state BIC mean =1.82e+04 ±137.52, CI [1.78e+04, 1.83e+04]). The BIC overlap of 95% confidence intervals between two-state and one-state was 1.98% for strong and 85.64% for weak learners. The smaller overlap of BIC intervals in the strong learners' group indicates they rely on the combination of fast and slow processes. In contrast, weak learners rely more on a single learning process. Error bars represent 95% confidence BIC interval.
